# Transcriptome analysis of alcohol dependence and stress interactions in the nucleus of the solitary tract

**DOI:** 10.1101/2022.03.17.484799

**Authors:** EK Grantham, GR Tiwari, O Ponomereva, AS Warden, A DaCosta, S Mason, YA Blednov, RA Harris, MF Lopez, HC Becker, RD Mayfield

## Abstract

Stress exposure contributes to the development of drug and alcohol use disorders. In animal models, stress exacerbates escalations in alcohol consumption in alcohol-dependent animals. The nucleus of the solitary tract (NTS) is a critical brainstem region for integrating and relaying peripheral signals to regulate stress responses. To define the molecular adaptions within this brain region that may contribute to stress-induced alcohol drinking, we exposed animals to chronic intermittent bouts of ethanol vapor (CIE), forced swim stress (FSS), or both (CIE + FSS) and then transcriptionally profiled the NTS at three different timepoints after the last vapor exposure (0-hr, 72-hr, and 186-hr). We identified interferon (IFN) signaling as a critical gene network correlated with alcohol consumption levels. Using a likelihood ratio test, we identified genes that were differentially expressed across time and between groups. Clustering analysis of these genes to identify unique expression patterns identified a subset of genes that fail to normalize in the CIE + FSS group, but not the others. These genes were enriched for cell-to-cell interaction and cellular movement pointing to long-term structural and functional changes in this brain region caused by the unique interaction of alcohol dependence and stress. Specific genes of interest identified in this group include *Aqp4*, *Il16*, *Reln*, *Grm4*, *Gabrd*, and *Gabra6*. We also compared gene expression changes in the NTS to the PFC and found a significant overlap of genes between the two brain regions. Overlapping NTS/PFC genes in the CIE + FSS group were enriched for type I IFN signaling. Finally, we tested the hypothesis that activation of type I IFN signaling increases alcohol consumption based on the three lines of evidence identifying type I IFN signaling as critical for escalations in alcohol intake. Mice treated with recombinant IFNβ showed significantly elevated levels of alcohol intake in a two-bottle choice procedure compared to saline-treated controls. Overall, these results define the transcriptomic changes across time in the NTS that may be critical to the development of stress-induced increases in alcohol consumption and alcohol dependence.

## Introduction

Stress is a major contributor to the development of substance use disorders, including alcohol use disorder (AUD)[1–3]. Chronic alcohol exposure and withdrawal perturb the hypothalamic-pituitary-adrenocortical (HPA) axis[3]. Repeated HPA axis perturbations by stress and alcohol exposures lead to molecular neuroadaptations that may mediate compulsive alcohol consumption and ultimately the development of AUD[4–6]. Therefore, it is critical to study the molecular processes affected by chronic alcohol exposure and stress in stress-regulating brain regions to understand underlying maladaptive behaviors that lead to AUD.

The nucleus of the solitary tract (NTS), a critical brainstem regulator of stress responses[4]. The NTS integrates peripheral visceral inputs with local brainstem circuits to coordinate autonomic and endocrine homeostatic functions[5–8]. The NTS is highly interconnected with stress-regulating circuits like the extended amygdala and hypothalamus and has been associated with anxiety and stress, positioning it as a critical brain region for the bottom-up control of stress-induced alcohol consumption[4,9–15].

Acute and chronic exposure to ethanol have been shown to activate the NTS as measured by c-Fos induction[16–19]. Activity within the NTS has been associated with substance use withdrawal[20]. Activation and inhibition of the NTS regulate both stress reactivity and alcohol consumption[4,12,13,21]. Despite the known role of the NTS in stress responses and evidence for its role in alcohol behaviors, it is unclear how the NTS regulates stress-induced alcohol consumption and molecular adaptations that occur in response to stress and combined chronic alcohol exposure have not been explored in the NTS.

In the present study we employed a model of alcohol drinking that involves stress exposure alone and/or in combination with chronic alcohol exposure. Specificallly, the model demonstrates that repeated brief forced swim stress (FSS) exposure interacts with chronic intermittent ethanol (CIE) exposure to enhance alcohol consumption in dependent (CIE-exposed) mice but FSS does not alter more moderate levels of alcohol intake in nondependent mice (Anderson et al., 2016; Hau et al., 2022; Lopez et al., 2016). We hypothesized that gene expression changes in the NTS in the CIE-FSS Drinking model would define the molecular drivers of stress-enhanced alcohol consumption. Our results identify neuroimmune signaling, specifically interferon (IFN) α/β signaling, as a critical biological pathway perturbed by CIE-alone and CIE+FSS conditions. We validated a functional role for IFNβ signaling by administering recombinant IFNβ during a voluntary drinking test. We also identified a network of genes with a unique expression pattern across time in the CIE+FSS group that suggests the interaction between CIE and FSS causes a unique transcriptional signature related to cell-to-cell signaling. Overall, our results highlight transcriptional networks related to unique interactions of stress and chronic alcohol exposure on voluntary alcohol consumption and offers new avenues for AUD therapeutic drug development.

## Methods

### Animal Model

Information about animals used for transcriptional analysis can be found in a previous publication[22]. In short, adult male C57BL/6J mice were used for transcriptional profiling after treatment in the CIE-FSS Drinking model (PMID: 8146231). In vivo exposure was conducted at MUSC and in accordance with Institutional Animal Care and Use Committee and NIH Guidelines The medial prefrontal cortex (mPFC) was isolated and transcriptionally profiled[22] and the nucleus of the solitary tract (NTS) was isolated and transcriptionally profiled separately for this publication.

### Procedure

Mice were treated in the CIE-FSS Drinking model involving chronic intermittent ethanol (CIE) vapor exposure and forced swim stress (FSS) exposure, as described above (Supplemental Fig. 1a) and described in detail in previous work[23,24]. Briefly, mice were first trained to drink 15% (v/v) ethanol with access for 1-hr/day starting 3-hr after lights off. Once stable baseline intake was established (~3 weeks), mice were separated into four groups (equated for baseline level of alcohol intake): CNTL, CIE-alone, FSS-alone, and CIE+FSS. Drinking tests were performed in between CIE or AIR (CNTL) exposure cycles using the same drinking procedure used for baseline measurements. Mice were decapitated after their fifth and last CIE or AIR (CNTL) exposure (0-hr, 72-hr, or 168-hr), and whole brains were immediately snap-frozen. Drinking data and blood ethanol concentrations for the animals used in this experiment can be found in previously published work[22].

### RNA Sequencing and Bioinformatics Analysis

Isolated total RNA from mouse NTS (Supplemental Fig.1b) was submitted to the Genomic Sequencing and Analysis Facility at The University of Texas at Austin. Sequencing libraries were constructed using a 3’ Tag-based approach (TagSeq), targeting the 3’ end of RNA fragments from ~16 ng/μL of each RNA sample. Samples were sequenced on the HiSeq 2500 (Illumina) platform with a read depth of approximately 7.6 million reads (single-end 100 bp reads; 4.6 million high quality reads per sample after trimming). A total of 89 samples were included with an average of 1.8 million uniquely mapped reads after mapping. TagSeq detected a total of 56,262 transcripts. On average, ~38,000 transcripts per sample were detected, representing ~22,000 protein coding genes. Read quality was assessed using MultiQC (version 1.7). Reads were mapped to the mouse reference genome (Gencode GRCm38.p6 release M25) using a STAR (version STAR_2.5.4b) aligner. Raw counts were quantified using HTSeq (version 0.11.2).

### Differential Gene Expression Analysis and Time-Course Cluster Analysis

The R (version 3.6.1) package DESeq2 (version 1.22.2) was used to identify differentially expressed genes (DEGs) between groups at each timepoint using the default Wald Test within groups or using a likelihood ratio test (LRT) across different levels for clustering analysis using the *DESeq* function. For comparisons within timepoints, nominal p<0.05 was selected to ascertain shared and nonshared changes in gene expression. For comparisons across timepoints and groups, the clustering function *degPatterns* from the R package DEGReports was used on regularized log transformation of the normalized counts of DEGs identified using LRT on the full DESeq model of *~Group* + *Time* + *Group:Time*, rather than the reduced model *~Group* + *Time*. *degPatterns* uses hierarchical clustering based on pair-wise correlations to identify groups of genes similarly expressed across time. This analysis produced clusters (*n* = 11) of genes with similar expression profiles across time and within groups. Statistically significant changes were limited to genes with FDR<0.05

### Weighted Gene Co-expression Network Analysis (WGCNA)

The R package WGCNA[25] (version 1.69) was used to identify genes correlated with behavioral traits of interest. The general framework for WGCNA analysis has been previously described[26]. Briefly, we constructed a signed adjacency matrix by calculating Pearson correlations for all pairs of genes. To emphasize strong correlations on an exponential scale, we raised the adjacency to power B, which was re-calculated for each timepoint comparison, so the resulting networks exhibited approximate scale-free topology (scale free topology fit = 0.85). To identify gene modules, all genes were hierarchically clustered based on connection strength determined using a topological overlap dissimilarity calculation. Resulting gene dendrograms were used for module detection using the dynamic tree cut method (minimum module size = 100). To determine module-trait relationships, Pearson correlations were calculated for module eigengene expression with CIE treatment status and BAC. Resulting p-values from module-trait correlations were adjusted for multiple comparisons using FDR<0.05.

### Ingenuity Pathway Analysis (IPA)

Cluster gene lists were entered into Ingenuity Pathway Analysis[27] (version 1-19-02) with log2 fold-change, p-value, and baseMean as inputs. We used a nominal p<0.05 to identify DEGs. IPA identified enrichment for canonical pathways and molecular function based on gene expression patterns submitted.

### Gene Ontology and Cell-type Enrichment Analysis

DEGs were analyzed for enrichment of canonical gene ontologies and molecular pathways using the bioinformatic tool and R package, Enrichr [28,29] (version 3.0). Reported ontological categories and pathways were limited to top six terms for each timepoint within each of the separate experimental groups (p<0.05). DEGs (*Group* + *Time*) were compared to cell-type marker genes within the community-curated single-cell RNA sequencing dataset, Panglao Database[30] to determine the overrepresentation of the major CNS cell-types using a Fisher’s exact test (p<0.05).

### Rank-rank hypergeometric overlap test (RRHO)

We used a rank-rank hypergeometric overlap test (RRHO) to compare patterns of gene regulation between the NTS and PFC from the same animals. RRHO identifies overlap between expression profiles in a threshold free manner to assess the degree and significance of overlap[31]. Full differential expression lists were ranked by the −log10(p-value) multiplied by the sign of the fold change from the differential gene expression analysis. The RRHO2 Bioconductor package (version 1.0) was used to evaluate the overlap of differential expression lists between PFC and NTS in the CIE + FSS group across timepoints.

Heatmaps visualize hypergeometric p-values for significant overlap between sets of genes, where smaller p-values correspond to warmer colors.

Every-other-day two-bottle-choice (EOD-2BC) drinking and interferon-β administration: Adult male C57BL/6j mice were individually housed in standard polycarbonate shoebox cages on a 12-hr light/dark cycle at the University of Texas at Austin with food (TMH 1800 5LL2 chow) and water *ad libitum*. All procedures were approved by the University of Texas Institutional Animal Care and Use Committee and adhered to NIH Guidelines (AAALAC accredited). Intermittent access to ethanol increases voluntary drinking in rodents[32–35].The two-bottle choice protocol was carried out as previously described[36]. Briefly, mice were offered a choice between 15% ethanol (v/v) or water for 24-hr sessions and water only was offered on alternating days. The quantity of ethanol consumed (g/kg body weight/24-hr) was calculated for each mouse, and these values were averaged across two drinking days for presentation. Recombinant interferon-β (IFNβ) (8324-MB) was purchased from R&D Systems. It was reconstituted it at 100 μg/mL in sterile PBS containing 0.1% BSA and injected (i.p.) at a dose of 2.5 μg/injection. IFNβ was injected every 5th day (1 day before every new round of drinking, which consisted of 2 drinking days).

## Results

### Overview of differential gene expression

We identified differentially expressed genes (DEGs) by comparing the CNTL (AIR) condition with each treatment group (CIE-alone, FSS-alone, and CIE+FSS). The number of DEGs with nominal p<0.05 varied between 412 and 2020 genes (Fig. 1). The largest overlap was between CIE-alone and CIE+FSS groups at 0-hr, with 782 common DEGs (Fig. 1a). The FSS-alone group showed the largest number of DEGs at the 72-hr timepoint, in contrast to CIE-alone and CIE+FSS conditions, which both showed the largest number of DEGs at 0-hr (Fig. 1a-b). Many alterations in gene expression were unique to each treatment group and timepoint, suggesting each treatment produces a specific transcriptional signature in the NTS.

**Figure 1.**
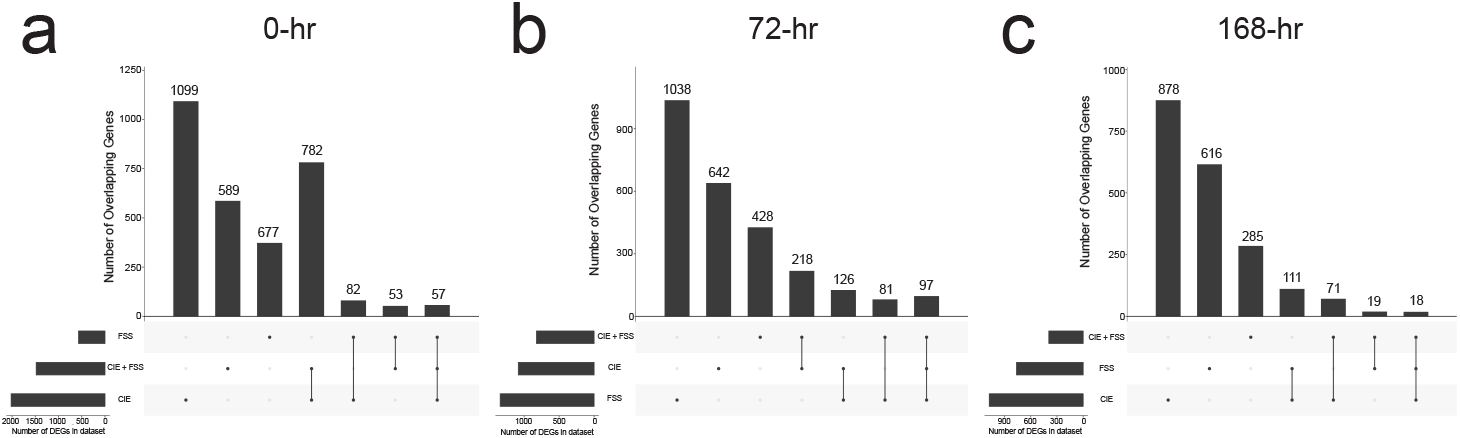
Number of differentially expressed genes by group and timepoint. The R package UpSetR (Version 1.4.0) was used to show the number of differentially expressed genes (DEGs) unique to each treatment group and common between treatment groups. Total number of DEGs per group shown in lower left corner of each plot. Filled in circles indicate which group(s) contribute to plotted number of DEGs above. a) Number of DEGs by group 0-hr timepoint. b) Number of DEGs by group 72-hr timepoint. c) Number of DEGs by group168-hr timepoint.

The three timepoints (0-hr, 72-hr, and 168-hr) allowed us to define transient, persistent, and compensatory gene expression patterns related to different stages of intoxication and withdrawal[37]. The 0-hr timepoint corresponds to immediate removal from the ethanol vapor inhalation chamber, when blood and brain alcohol levels are relatively high (e.g., 175-225 mg/dL). The 72-hr timepoint is considered to reflect the early protracted phase of withdrawal, when tremor and convulsions have substantially subsided but measures of negative affect (i.e., anxiety) may emerge (Heilig et al., 2010). The 72-hr timepoint is also critical because this is the timepoint at which animals resume access to their next drinking session, so the mice are presumably in a highly anticipatory state. The 168-hr timepoint is useful for evaluating whether persistent gene expression changes can be observed after protracted abstinence.

We sought to identify gene expression changes unique to the CIE+FSS group as they may point to biological neuroadaptations that drive stress-enhanced alcohol consumption. We identified 589, 429, and 291 DEGs unique to the CIE+FSS group across each time point, respectively (Fig. 1). Unique CIE+FSS DEGs at the 0-hr timepoint were enriched for glutamate signaling and regulation of glial cell differentiation (Table 2). Unique CIE+FSS DEGs at the 72-hr timepoint were enriched for steroid metabolic processes and cholesterol esterification (Table 2). Unique CIE+FSS DEGs at the 168-hr timepoint were enriched for death-inducing signaling complex and tyrosine kinase signaling (Table 2). These results suggest early neuroadaptations that involve synaptic and glial cell function and long-term changes in cell survival and intracellular signaling.

**Table 1.**
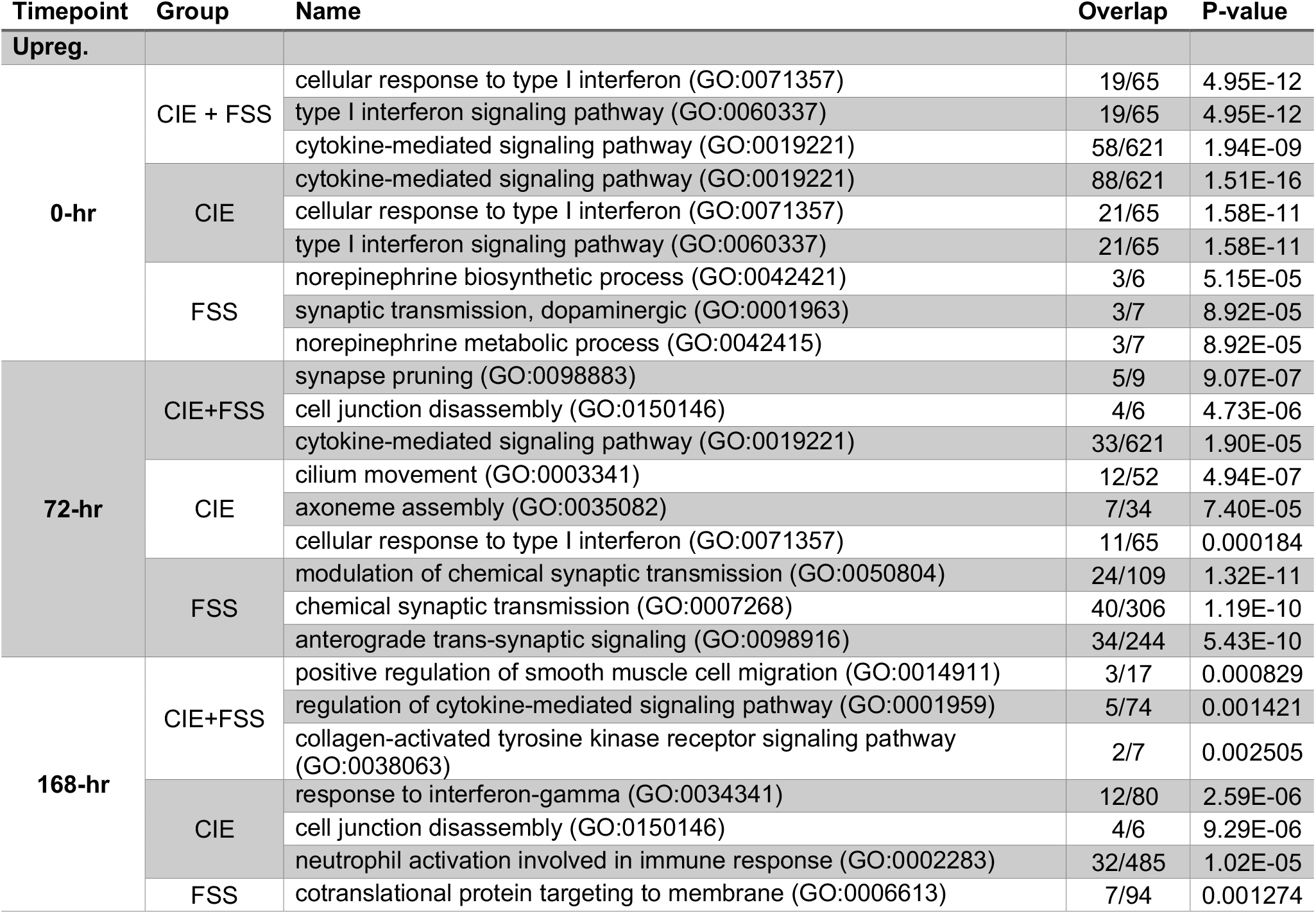
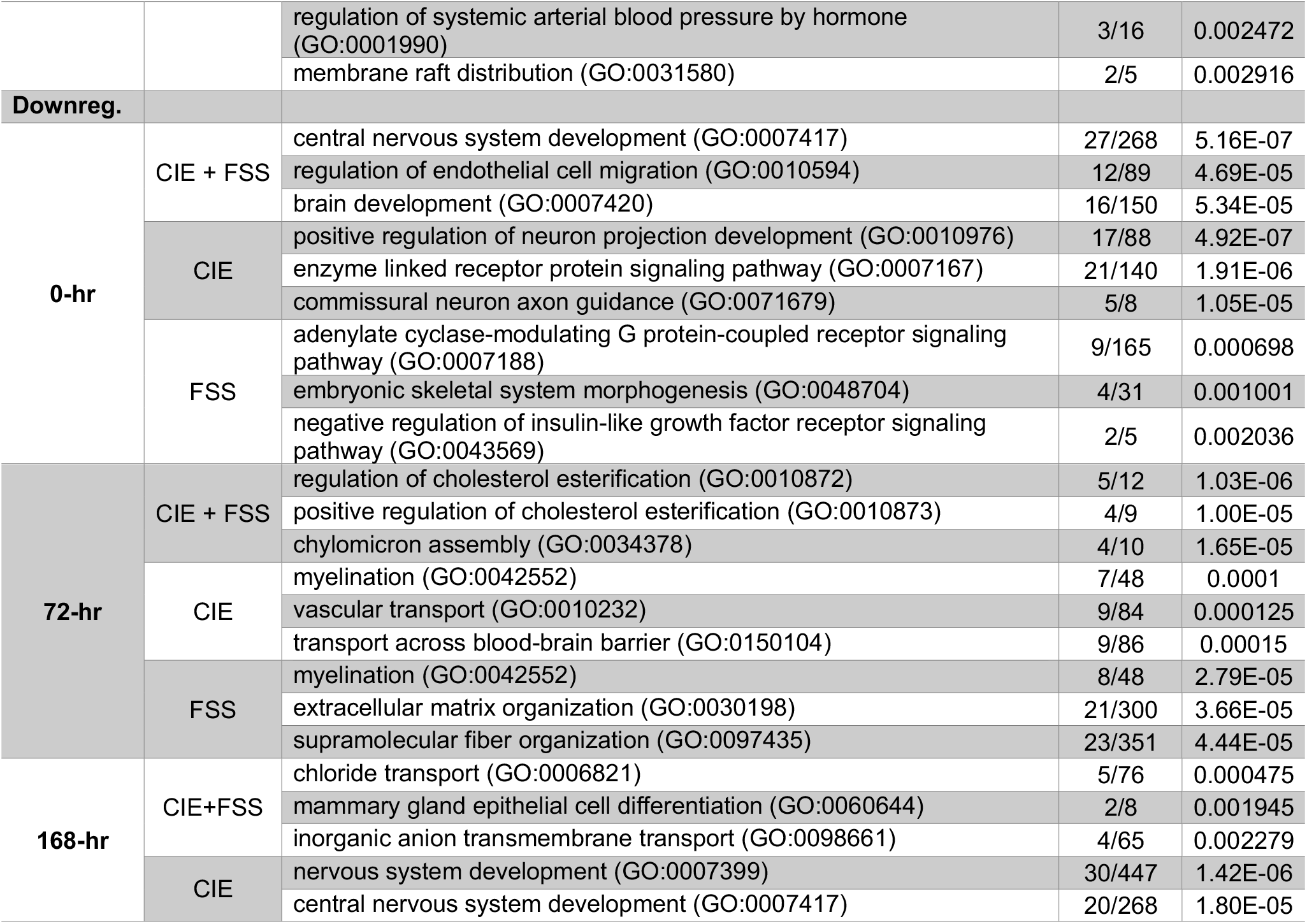
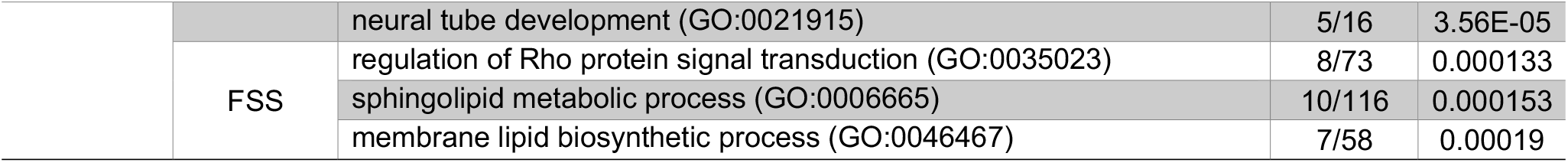
Enrichment of DEGs from the NTS by treatment group and timepoint, separated by upregulated (top) and downregulated (bottom) DEGs.

**Table 2.** Biological pathways uniquely differentially regulated by CIE + FSS in the NTS

| Timepoint | Name | Overlap | P-value |
| --- | --- | --- | --- |
| 0-hr | glutamate receptor signaling pathway (GO:0007215) | 7/37 | 8.8491E-05 |
|  | regulation of glial cell differentiation (GO:0045685) | 3/7 | 0.00081403 |
|  | vascular associated smooth muscle cell differentiation (GO:0035886) | 3/8 | 0.001274 |
| 72-hr | positive regulation of steroid metabolic process (GO:0045940) | 5/13 | 2.23E-40 |
|  | positive regulation of cholesterol esterification (GO:0010873) | 4/10 | 2.84E-38 |
|  | leukocyte tethering or rolling (GO:0050901) | 5/18 | 3.09E-31 |
| 168-hr | collagen-activated tyrosine kinase receptor signaling pathway (GO:0038063) | 2/7 | 0.00405324 |
|  | death-inducing signaling complex assembly (GO:0071550) | 2/7 | 0.00405324 |
|  | mammary gland epithelial cell differentiation (GO:0060644) | 2/8 | 0.00535363 |

We also identified DEGs in the FSS-alone treatment group (Fig. 1). Although these animals do not show an increase in drinking, these data are useful for understanding the transcriptional changes caused by chronic repeated stress in a critical stress-regulating brain region and may inform other behavioral phenotypes of interest. For example, repeated FSS has been shown to alter performance in learning and memory behavioral tasks[38,39]. Overall, early and intermediate transcriptional changes in this group point to significant alterations in synaptic function in the NTS after FSS exposure (Table 1). Given the abundance of changes in neuroimmune genes observed in both the CIE-alone and CIE+FSS groups, we decided to specifically evaluate differential expression of neuroimmune genes in the FSS-alone group. We identified a pattern where neuroimmune genes that were upregulated in the CIE-alone and CIE+FSS group were at some point downregulated in the FSS-alone group: *Ccl8*, *Il10ra*, *Il1r1*, *Ifi27*, *Il16*,and *Tnfrsf1a*. Importantly, this gene expression pattern does not translate to an alcohol drinking phenotype but may lead to susceptibility to excessive drinking when combined with chronic ethanol exposure.

Genes tend to work in coordination, referred to as gene networks, across time and in response to different manipulations. To identify gene networks related to excessive alcohol intake and treatment groups, we used Weighted Gene Co-expression Network Analysis (WGCNA). We identified several modules in each timepoint that correlated negatively or positively with alcohol intake on the last test day (Test 4), blood ethanol concentration (BEC), and change in alcohol intake.

### Immune regulatory gene networks are positively correlated with alcohol consumption at 0-hr and 72-hr

At the 0-hr timepoint, the turquoise module was positively correlated with alcohol intake on the last test day, while the brown, tan, and red modules were negatively correlated with alcohol intake (Supplemental Fig. 2). Turquoise eigengene expression was upregulated in both CIE-alone and CIE+FSS groups (Fig. 2b), consistent with the idea that these genes are positively correlated with alcohol consumption on the final drinking session (Test 4); these treatment groups consumed the highest levels of alcohol relative to the FSS-alone and CNTL groups[22]. The turquoise module was enriched for interferon (IFN) α/β signaling, immune signaling by interferons, interleukins, prolactin, and growth hormones, and antigen processing (Fig. 2c). All hub genes (genes with highest number of connections within the network) in the turquoise module were also significantly upregulated in the CIE+FSS group relative to control (Fig 3d). Overall, the turquoise module represents a gene network that is significantly perturbed by and correlated with increased alcohol consumption associated with stress-alcohol interactions.

**Figure 2.**
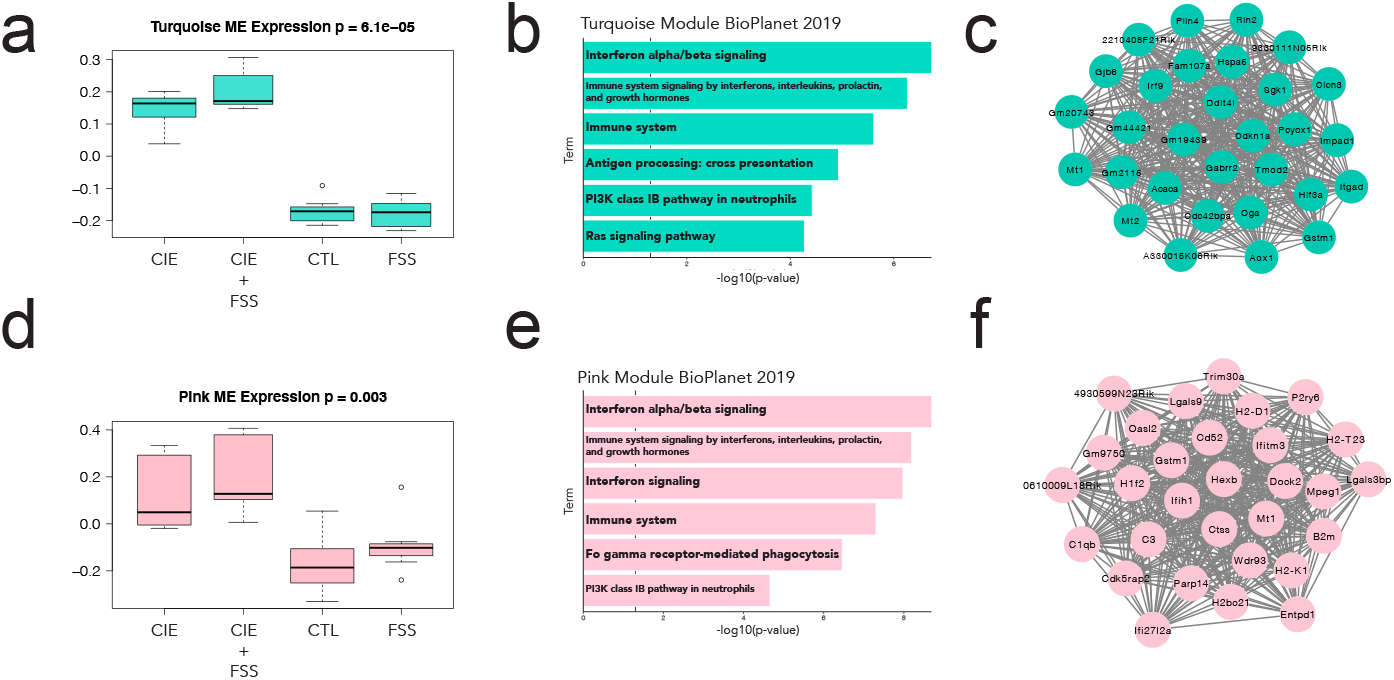
Weighted Gene Co-expression Network Analysis (WGCNA) modules correlated with alcohol intake at 0-hr and 72-hr. a) Turquoise module eigengene expression by group. b) Enrichment of turquoise module using BioPlanet repository 2019. c) Top 30 hub genes for turquoise module. d) Pink module eigengene expression by group. e) Enrichment of pink module using BioPlanet repository 2019. f) Top 30 hub genes for pink module.

**Figure 3.**
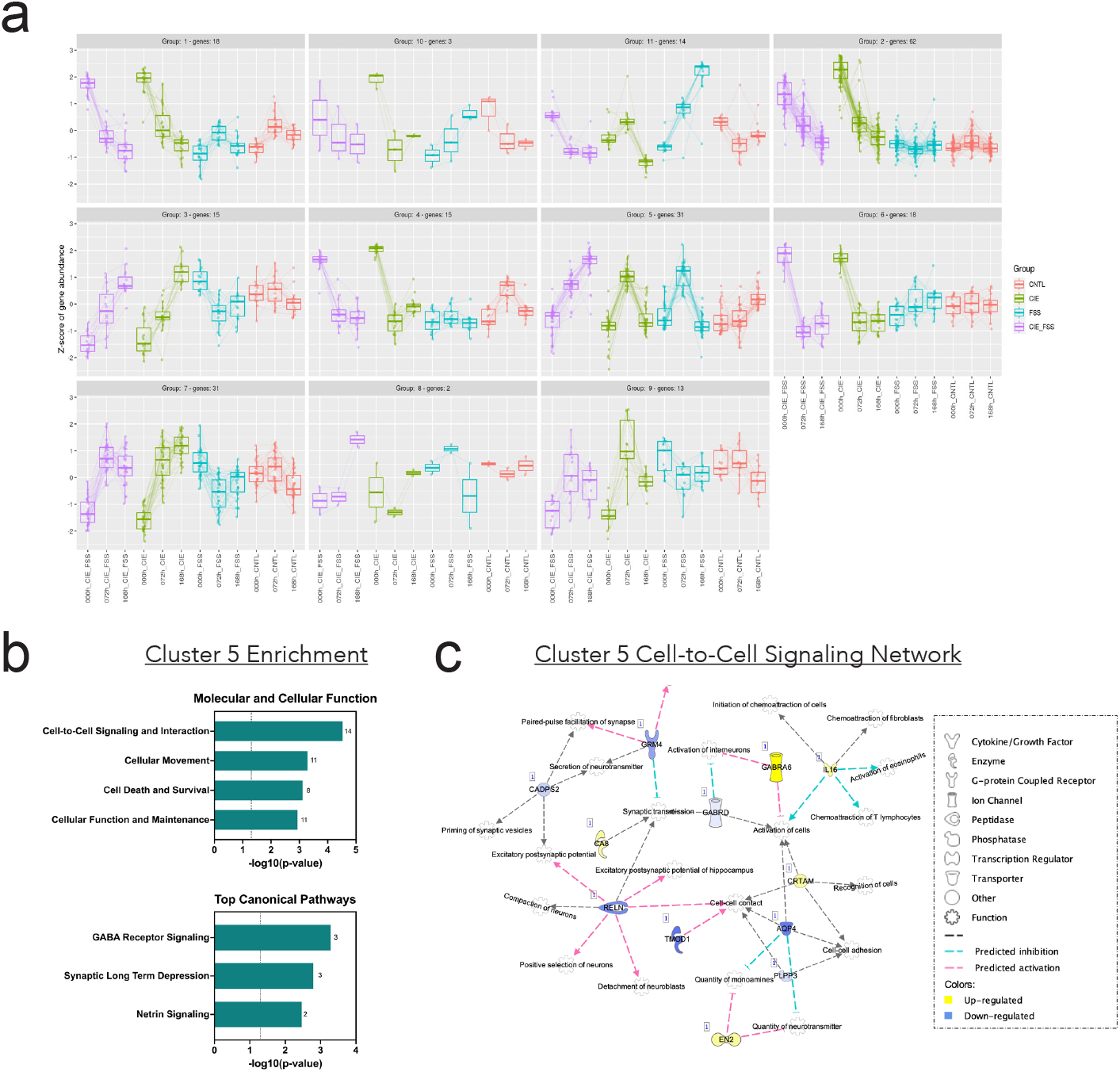
Clustering analysis of significantly differentially expressed genes within groups across time. a) Cluster plots with clusters of uniquely expressed genes (DEGs) across time within groups plotted against the Z-score of gene abundance. b) Enrichment analysis of Cluster 5: Molecular and Cellular Function and Top Canonical Pathways using IPA. c) Cell-to-Cell Signaling and Interaction Network visualized with IPA software.

At the 72-hr timepoint, the only module positively correlated with alcohol consumption was the pink module (r^2^ = 0.59, p = 0.002)(Supplemental Fig. 2). The pink module eigengene expression was upregulated in both CIE-alone and CIE+FSS groups (Fig. 2d). Similar to the turquoise module at the 0-hr timepoint, the pink module was enriched for IFNα/β signaling and neuroimmune signaling such as interleukin, prolactin, growth hormone and type II IFN signaling (Fig. 2e). The top 30 hub genes from the pink module include several genes that have been previously studied in the context of alcohol use disorder and voluntary alcohol consumption, including *B2m* and *Ctss* (Fig. 2f).

### Clustering analysis identifies a unique subset of genes related to cell-to-cell signaling in the CIE + FSS group

While WGCNA is useful for identifying networks of genes correlated with a trait of interest, like alcohol consumption, none of the modules identified were uniquely correlated with the CIE+FSS group. This is likely because gene expression changes that are unique to this group are smaller in scale than what WGCNA can detect (minimum module size = 100 genes). Evaluating gene expression changes at three different time-points allows for analysis of temporal patterns of expression that may be unique to a particular treatment group. To identify gene expression signatures unique to the experimental CIE+FSS group, we used *degPatterns* function from the R package DEGreports (cite package again). We identified 224 significantly DEGs using the likelihood ratio test (LRT) with an adjusted p< 0.05. The function *degPatterns* then clustered these significant gene expression patterns into 11 different clusters with similar expression profiles (Fig. 3a).

The CIE-alone and CIE+FSS DEGs typically showed very similar expression patterns across time, except for one cluster: 5. In this cluster, gene expression in the CIE+FSS group increased stepwise across the three timepoints, whereas, the same genes showed more transient changes in the other treatment groups. These genes were enriched for cell-to-cell signaling, cellular movement, and GABA receptor signaling (Fig. 3b), suggesting long-term structural and functional changes in the NTS that are unique to the interaction of stress and chronic alcohol exposuyre (CIE+FSS condition). Upon closer inspection of the cell-to-cell signaling network, which had the largest number of DEGs in cluster 5, many of these genes are related to immune function (*Il16*), synaptic signaling (*Gabra6*, *Gabrd*, *Ca8*, *Grm4*) and cell-to-cell contact (*Reln*, *Aqp4*, *Tmod1*) (Fig. 3c).

### Overlapping expression profiles between the NTS and PFC

Previous work with the same animals identified transcriptional changes associated with CIE+FSS in the mouse cortex[22]. Cortical transcriptional signatures from the same alcohol-dependent, stressed animals provide an opportunity to identify overlapping gene expression changes between PFC and NTS. Tracing studies show that the medial prefrontal cortex (mPFC) sends direct projections to the NTS[40] and the mPFC may act as a depressor of NTS activity[41]. Therefore, we sought to identify correlated gene expression patterns between the NTS and mPFC from alcohol-dependent, stressed animals using rank-rank hypergeometric overlap tests (RRHO)[31,42]. The RRHO algorithm assesses the overlap in DEGs between PFC and NTS in the CIE+FSS group to identify concordant and discordant patterns in gene expression between two datasets (i.e., genes that are co- up- or down-regulated or oppositely up- or down-regulated between the two brain regions).

The largest observed overlap was at the 0-hr timepoint (max −log10(p-value) = 136) in genes upregulated in PFC and NTS (Fig. 4a). There were 253 genes identified as significantly differentially expressed in both PFC and NTS (Fig. 4b). Enrichment analysis identified a strong type I IFN signaling signature (p = 2.88 × 10^-11^) (Fig. 4b). There was a weaker overlap in genes co-downregulated at the 0-hr timepoint (max −log10(p-value) = 55) (Fig. 4a), suggesting that CIE+FSS in general leads to more common upregulated DEGs across brain regions compared to downregulated DEGs. There were 142 common DEGs in both PFC and NTS (Fig. 4c). These core-overlapping genes were enriched for regulation of long-term synaptic depression and regulation of microtubule-based process (Fig. 4c), suggesting synaptic and structural changes to both of these brain regions caused by the combination of chronic alcohol (CIE) and stress (FSS) experience.

**Figure 4.**
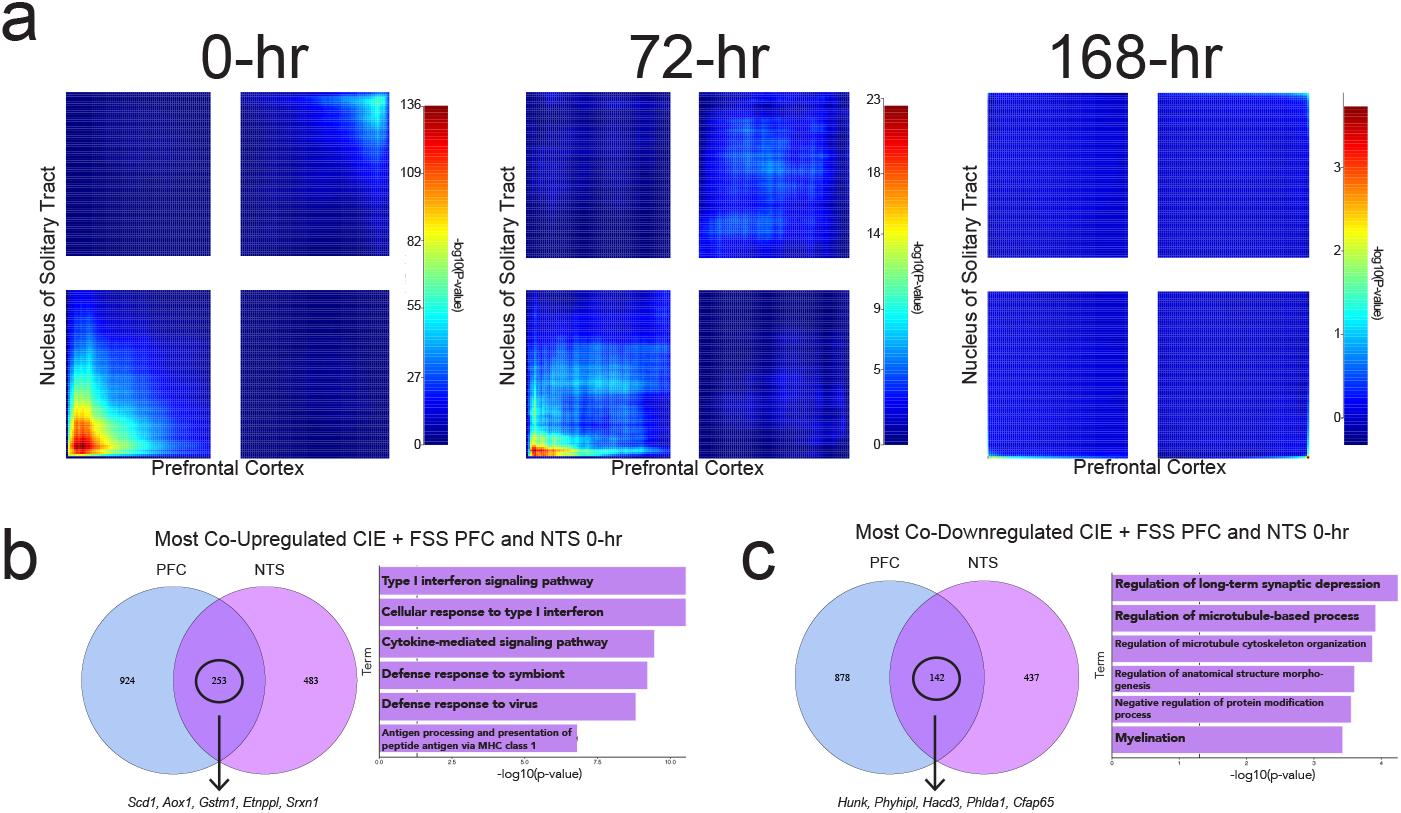
Overlapping differentially expressed genes in CIE + FSS group between NTS and PFC. a) RRHO plots of overlapping differentially expressed genes at 0-hr, 72-hr, and 168-hr between NTS and PFC CIE+FSS groups. b) Venn diagram of coordinately upregulated genes from the lower left quadrant of 0-hr RRHO plot (left). Arrow points to top 5 overlapping genes in rank-rank gene list. Enrichment of 253 coordinately upregulated genes in NTS and PFC (right). c) Venn diagram of coordinately downregulated genes from the upper right quadrant of 0-hr RRHO plot (left). Arrow points to top 5 overlapping genes in rank-rank gene list. Enrichment of 142 coordinately downregulated genes in NTS and PFC (right).

Co-upregulated genes also showed a significant overlap at the 72-hr timepoint (max −log10(p-value) = 23) (Fig. 4a), though to a lesser degree than the 0-hr timepoint. At the 168-hr timepoint, there was no significant overlap for co-regulated or discordantly regulated genes (Fig. 4a). These results show that the CIE+FSS treatment produces similar expression patterns across two distinct brain regions at early and intermediate timepoints, but not after protracted withdrawal.

### Administration of IFNβ demonstrates causal role for type I interferon signaling in voluntary alcohol consumption

Type I IFN signaling was identified as a potential regulator of voluntary alcohol consumption using differential gene expression analysis, WGCNA, and RRHO tests between NTS and PFC. Based on these results, we hypothesized that perturbation of type I IFN signaling would impact voluntary alcohol consumption, regardless of stress exposure. Specifically, increased expression of type I IFN signaling components positively correlated with voluntary alcohol intake levels. Thus, we tested the hypothesis that activation of type I IFN signaling would increase voluntary alcohol intake using an every-other-day two-bottle-choice (EOD-2BC) drinking procedure. Systemic administration of the type I IFN, IFNβ. IFNβ (2.5 μg) treatment increased voluntary alcohol consumption and preference (Fig. 5a-b), with no effect on total fluid intake (Fig. 5c). These results support our hypothesis that activation of type I IFN signaling drives increases in voluntary alcohol consumption.

**Figure 5.**
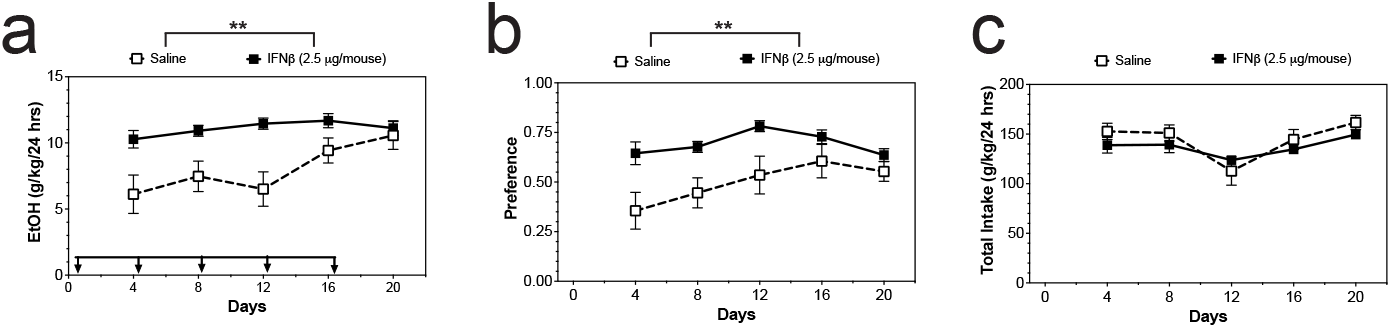
Activation of type I interferon (IFN) signaling increases voluntary alcohol consumption. a) C57BL/6J mice were injected with recombinant IFNβ was injected on every 5^th^ day during EOD drinking. IFNβ increased alcohol intake, measured in g/kg body weight averaged across two drinking days. b) IFNβ increased alcohol preference, measured in the ratio of ethanol to water consumed and averaged across two drinking days. c) IFNβ had no effect on total fluid intake. (*p<0.05, **p<0.01, ***p<0.001, n = 9-10/group)

## Discussion

In this paper, we identify molecular networks associated with alcohol dependence and stress using a systems genomics approach that integrates co-expression network analysis, temporal clustering analysis, and behavioral analysis. Starting with differential gene expression analysis, we identified a consistent upregulation of type I IFN signaling in alcohol-dependent animals. For example, DEGs in both CIE-alone and CIE+FSS groups at 0-hr and 72-hr were enriched for type I IFN signaling and immune signaling. Neuroimmune signaling has been identified as a major biological system perturbed by chronic alcohol in both animal models and post-mortem human alcoholic brain tissue (reviewed [43]). The identification of neuroimmune-related genes in the NTS suggests neuroimmune signaling is potentially perturbed in a coordinated manner within several brain regions important for the pathology of alcohol use disorder (AUD).

Across time, there were very few persistent gene expression changes, regardless of treatment group. Three genes, *Ccl5*, *Ly86*, and *Ttyh1*, were differentially expressed across all three timepoints in both the CIE-alone and CIE+FSS groups. Only *Ccl5* was upregulated across all three timepoints in both groups, suggesting long-term alterations in transcriptional regulation of this gene. *Ccl5* is involved in inducing the migration of phagocytes across the blood brain barrier to sites of inflammation[44] and has been shown to be involved in the recruitment of peripheral macrophages into the CNS in response to chronic alcohol consumption[45]. The persistent upregulation of *Ccl5* and its known involvement in alcohol-induced neuroimmune activation make it a promising target for future study in the context of AUD.

Co-expression network analysis was useful for identifying gene networks correlated with alcohol intake and therefore may play a causal role in the regulation of excessive alcohol consumption[25,46–48]. Genes of interest within the 0-hr turquoise hub gene network include neuronal genes *Clcn3* and *Gabrr2*, glucocorticoid-inducible genes *Mt1*, *Mt2*, and *Sgk1*, immune-related genes *Irf9*, *Itgad*, and *Gstm1*, and extracellular-matrix-related genes *Tmod2*, *Gjb6*, and *Plin4*. The GABA-A receptor subunit rho2 (*Gabrr2*) is inhibited by ethanol and has been genetically linked to alcohol behaviors in mice and human alcohol dependence[49]. *Clcn3* is upregulated in in the nucleus accumbens of a genetic rodent model of excessive alcohol consumption[50]. In the brain, metallothioneins 1 and 2 (*Mt1* and *Mt2*) are glucocorticoid-inducible astrocytic proteins that have been shown to have a neuroprotective role[51–56]. Metallothioneins have been relatively understudied in the context of AUD, however *Mt2* brain expression has been associated with ethanol preference in mice[57]. Similarly, glucocorticoid-regulated kinase (*Sgk1*) blood levels are regulated by ethanol[58]. Finally, the extracellular matrix is a critical cellular component that appears to be perturbed by ethanol[59–61], warranting future study of the ECM-related genes identified in the turquoise network.

Several hub genes within the 72-hr pink module were similarly associated with neuroimmune signaling and have been implicated previously in alcohol research. For example, *B2m* and *Ctss* knockdown reduce alcohol consumption in a mouse model[62]. Several hub genes are also microglia-specific and have been implicated in the role of microglial-mediated escalations in alcohol consumption, including *C1qb*, *C3*, and *Ifitm3* [63–66]. Of note, *Mt1* and *Gstm1* were also hub genes in the turquoise module, suggesting expression of these genes in the NTS may be critical for escalations in alcohol consumption. Glutathione-S transferase mu 1 (*Gstm1*) promotes astrocyte-driven microglial activation during brain inflammation[67]; prolonged dysregulation of this gene in the NTS may be important for microglial-mediated escalations in alcohol-dependent animals[63].

Temporal clustering analysis helped identify a group of dysregulated genes unique to the CIE+FSS condition. Notably, none of the genes identified in this cluster were similarly identified as dysregulated in the PFC. There were several genes within this cluster that have previously been implicated in alcohol research, including *Gabra6* and *Gabrd*. *Gabra6* has been genetically linked to human alcohol dependence[68–70] and variations in human stress response[71,72]. Indeed, the GABAergic system is widely known as a critical regulator of stress responses and is a common drug target for the treatment of stress-related psychiatric conditions (depression, anxiety)[73]. *Gabra6* and *Gabrd* were also both upregulated in the brains of alcohol-drinking P rats[74]. Expression of these components of the GABAergic system in the NTS may play an important role in the regulation of stress-induced escalations in alcohol consumption. [75]

While the overarching aim of this study was to identify mediators of stress-induced excessive alcohol consumption, we were also able to identify a potential target for dependence-induced elevations in alcohol consumption using the two CIE-treated groups. A clear pattern emerged across all analyses pointing to type I IFN signaling as a mediator of dependence-induced escalations in alcohol consumption. We tested the hypothesis that activation of type I IFN signaling drives increases in alcohol consumption using recombinant IFNβ during EOD-2BC drinking. The main limitation of this causal link is the difference in behavioral model of drinking between the CIE procedure and EOD-2BC. EOD-2BC models similar aspects of AUD compared to CIE, including escalation of drinking and dependence[76]. EOD-2BC and CIE also show similar transcriptional profiles in PFC, suggesting that the molecular mechanisms of escalations in drinking are similar between the two models[48,77]. Therefore, we used the more time- and cost-effective EOD-2BC to test our functional hypothesis that type I IFN signaling mediates voluntary alcohol consumption.

Overall, this work adds to and extends the body of literature implicating neuroimmune signaling as a critical biological pathway involved in excessive drinking phenotypes. To our knowledge, our results are the first to demonstrate a role for neuroimmune signaling genes in stress-enhanced drinking. Further, these results provide new insights on transcriptional changes in a critical stress-regulating brain region that has thus far been understudied in the alcohol research field (NTS). We identified a network of genes unique to the CIE+FSS condition that may play a role in stress-induced escalations in alcohol consumption, and we identified a causal role for IFNβ in voluntary alcohol consumption.

## Notes

### Competing Interest Statement

The authors have declared no competing interest.

