## Supplemental Figures for "Transcriptome analysis of alcohol dependence and stress interactions in the nucleus of the solitary tract"

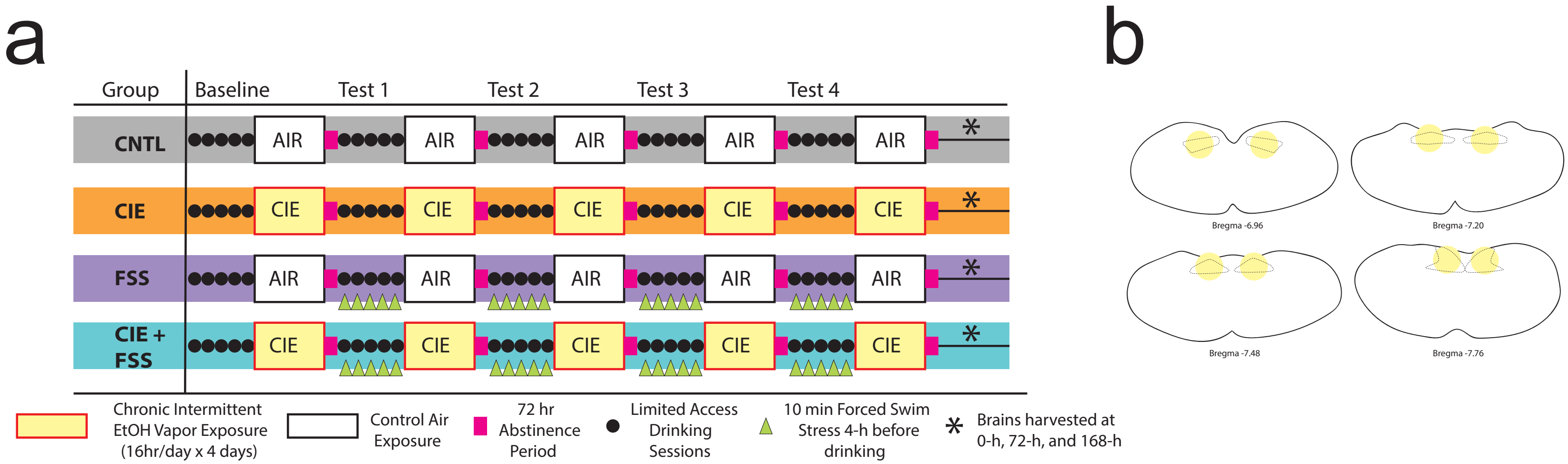


**Supplemental Figure 1. Experimental Design.**

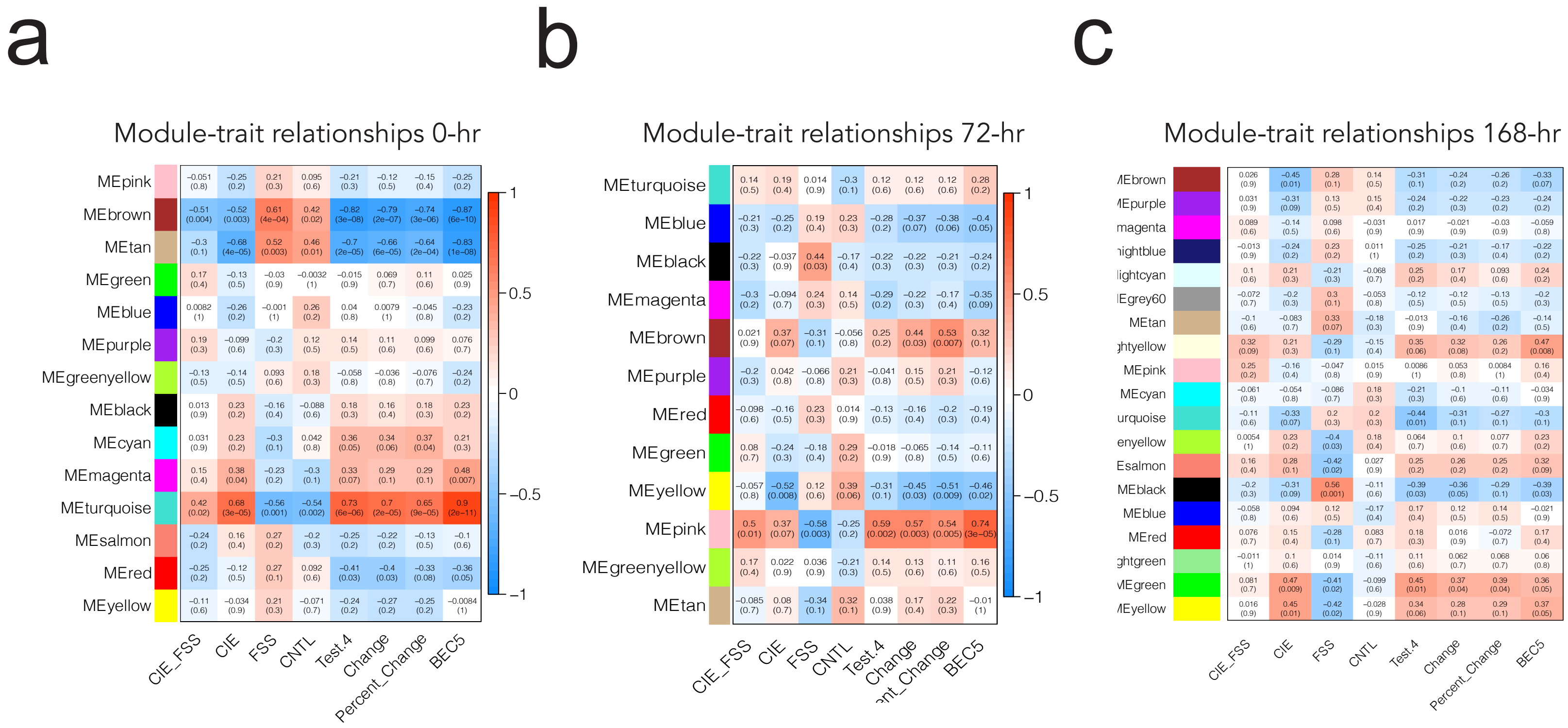


**Supplemental Figure 2. Module-trait relationships at 0-hr, 72-hr, and 168-hr.**
